# Direct colorimetric LAMP assay for in-field detection of African swine fever virus: a validation study during an outbreak in Vietnam

**DOI:** 10.1101/2020.06.03.132944

**Authors:** Diem Hong Tran, Hau Thi Tran, Uyen Phuong Le, Xuan Dang Vu, Thi Bich Ngoc Trinh, Hoang Dang Khoa Do, Van Thai Than, Le Minh Bui, Van Van Vu, Thi Lan Nguyen, Huong Thi Thu Phung, Van Phan Le

## Abstract

African Swine Fever (ASF) is a highly infectious viral disease with high mortality. The most recent ASF outbreak in Vietnam occurred in 2019, posing a threat to spread to the neighboring Asian countries. Without a commercial vaccine or efficient chemotherapeutics successfully developed, rapid diagnosis and necessary biosecurity procedures are required to control the disease. While the diagnosis method of ASF recommended by the World Organization of Animal Health is real-time PCR, it is not suitable for in-field detection of the disease. In this study, a colorimetric Loop-Mediated Isothermal Amplification (LAMP) assay was developed and evaluated for ASF virus detection using crude serum samples collected from domestic pigs in Vietnam during the 2019 outbreak. The LAMP results can be readily visualized to naked eyes within 30 minutes without the requirement of DNA extraction and sophisticated equipment. The sensitivity, specificity, and limit of detection of colorimetric LAMP assay were comparable to a commercial diagnostic real-time PCR kit. Results strongly indicate that the developed colorimetric LAMP assay is highly recommended for the in-field diagnosis of ASF.

## INTRODUCTION

African swine fever (ASF) is a contagious and dangerous viral disease of swine with extremely high mortality; in some cases up to 100% infected pigs were died (Galindo & Alonso, 2017; Penrith & Vosloo, 2009). ASF is caused by the ASF virus (ASFV), which is the only known member of the Asfivirus genus that belongs to the Asfarviridae family (Montgomery, 1921). ASFV is a large icosahedral virus that contains variable-length double-stranded DNA (ranging from 170 Kbp to 190 Kbp). After the first report in Kenya (Africa) in 1921, ASFV caused a transcontinental outbreak in Europe and South America in the middle of the 20^th^ century. In 2007, the virus was discovered in Georgia and continued to spread throughout the Eastern territories of the European Union in 2014 (Galindo & Alonso, 2017). In August 2018, China reported the first appearance of ASFV (Zhou et al., 2018). Not long after that Vietnam officially confirmed the first detection of ASFV in pigs in February 1^st^, 2019. The isolated viruses were 100% identical to those found in China and Georgia (Van Phan Le et al., 2019). Although ASFV does not directly threaten public health, it causes serious socio-economic interruption of the pig market, especially in countries where pork is considered as a predominant source of protein (Sánchez□Vizcaíno, Laddomada, & Arias, 2019).

Diagnosis of ASF is to identify the pigs that are infected with ASFV. The rapid and accurate detection of ASFV is important to distinguish those infected pigs from the others disease with similar clinical symptoms and thus, is required for timely control measures to prevent the disease from widely spread. The laboratory diagnosis of ASF includes the detection of ASFV-specific antigens and antibodies or viral genomic-DNA in clinical samples via virus isolation, serological or molecular assays. Accordingly, ASF diagnostic approach is divided into two groups: virological and serological (Cubillos et al., 2013; Oura, Edwards, & Batten, 2013). Many virological tests are currently available (Oura et al., 2013) involving virus isolation to detect the live ASFV (Malmquist & Hay, 1960), antigen ELISA (Vidal et al., 1997; Wardley, Abu Elzein, Crowther, & Wilkinson, 1979) or a fluorescent antibody test ELISA (Bool, Ordas, & Sanchez-Botija, 1969) to define viral antigens, polymerase chain reaction (PCR) assay (Agüero et al., 2003; Fernández□Pinero et al., 2013; King et al., 2003; McKillen et al., 2007; Steiger, Ackermann, Mettraux, & Kihm, 1992) and isothermal nucleic acid amplification tests (Frączyk, Woźniakowski, Kowalczyk, Niemczuk, & Pejsak, 2016; Hjertner, Meehan, McKillen, McNeilly, & Belák, 2005; James et al., 2010; Wang et al., 2020; Woźniakowski et al., 2017; Wu et al., 2016) to identify the viral genome. Meanwhile, the serological test (Cubillos et al., 2013) is based on the detection of specific antibodies utilizing the native or recombinant antigens (Alcaraz et al., 1995; C. Gallardo, Blanco, Rodríguez, Carrascosa, & Sanchez-Vizcaino, 2006; Pan, De Boer, & Hess, 1972; Pan, Huang, & Hess, 1982; Pastor, Laviada, Sanchez-Vizcaino, & Escribano, 1989; Perez-Filgueira et al., 2006). Serological methods were shown to have a limited sensitivity when the host antibody was present at low amount in the first few days of infection (M. C. Gallardo et al., 2015). On the contrary, PCR is the preferred diagnostic indicator of ASFV at the early stage of infection (M. C. Gallardo et al., 2015). The validated PCR assay could detect the viral DNA even when the clinical symptoms are not yet to be manifested (Beltran-Alcrudo, Arias, & Gallardo, 2017).

Currently, a quantitative real-time PCR method has been widely considered as one of the most precise techniques for virological diagnosis (Oura et al., 2013). Real-time PCR provides high accuracy, specificity and sensitivity and thus, has been recommended by the World Organization for Animal Health (OIE) for the detection of ASFV (World Organization for Animal Health, 2019). However, it requires cumbersome specialized equipment for a strict thermal-cycle control or result reading, as well as well-trained personnel, making it unsuitable for fast detection in the field. To overcome this drawback, different isothermal nucleic acid amplification methods, which are rapid and remarkably sensitive, have been established (Gill & Ghaemi, 2008). Unlike typical PCR, the isothermal nucleic acid amplification approach does not require changes in temperature and can be conducted without a need of a thermocycler (Craw & Balachandran, 2012). Loop-Mediated Isothermal Amplification (LAMP) has begun to gain huge attention since its first introduction (Notomi et al., 2000). LAMP requires just one kind of DNA polymerase without the need for modified or labeled DNA probes, making the experimental procedure less complicated and remarkably reducing the expense (Li, Li, Jia, & Yan, 2011). LAMP is more stable compared to PCR and real-time PCR under a wider range of temperature, pH and elongation time (Dhama et al., 2014; Francois et al., 2011). Moreover, LAMP was shown to have remarkable sensitivity and specificity with an extremely low limit of detection (LOD) compared to conventional PCR (Dhama et al., 2014; Zhang, Lowe, & Gooding, 2014). The presence of non-cognate DNA molecules in samples does not interfere with the sensitivity of LAMP (Iseki et al., 2007; Notomi et al., 2000; Zhang et al., 2014). Besides, LAMP is more tolerant to various PCR inhibitors such as trace quantities of whole-blood, hemin, urine or stools (Dhama et al., 2014; Francois et al., 2011; Kaneko, Kawana, Fukushima, & Suzutani, 2007). All those advantages make LAMP an excellent isothermal nucleic acid amplification technique which has been broadly utilized currently. Besides, to supply stable heat for amplification reactions, numerous electricity-free and portable devices were developed, allowing the simpler application of LAMP at field usage (Buser et al., 2015; Shah et al., 2015; Singleton et al., 2014; Singleton et al., 2013). In this study, we developed a rapid test kit based on colorimetric LAMP assay for the detection of ASFV utilizing a pH-sensitive dye for readout visualization. The test kit was extensively evaluated to detect ASFV using direct serum samples for an efficient in-field diagnosis.

## MATERIALS AND METHODS

### Sample collection and preparation

Blood and serum samples of the ASF symptomatic pigs from domestic pig farms in Vietnam were collected from October 2019 to January 2020. Viral genomic DNA was extracted from whole blood and serum samples using the QIAamp DNA Mini Kit (Qiagen, USA). Genomic DNA of ASFV was identified by a commercial real-time PCR kit (Median Diagnostics Inc., http://www.mediandiagnostics.com/asfv.php). The sample information and real-time PCR results are presented in Table S1.

### DNA preparation

The fragment (148500-148799) of Topoisomerase II gene of ASFV (GenBank accession No. AY261361.1) was obtained commercially (IDT, Singapore) (James et al., 2010). The synthetic DNA was used as the template for optimization and for identification of LOD of the LAMP assay. The sequence of LAMP primers was as described in the previous study (James et al., 2010) and the primer set was purchased from IDT (Singapore).

### PCR assay

The PCR was carried out in a 20 μl reaction volume containing 0.2 μM each of primers F3 and B3, 1 μl of DNA template, 0.2 μl of MyTaq DNA polymerase (Bioline, London, UK) and 4 μl of 5X MyTaq reaction buffer (Bioline, London, UK). The amplification products were analyzed by electrophoresis using a 2% agarose gel.

### ASF colorimetric LAMP reactions

WarmStart^®^ Colorimetric LAMP 2X Master Mix (DNA & RNA) was purchased from NEB (MA, USA). The LAMP assay reaction volume was 15 μl, consisting of 0.8 μM each outer primer (FIP and BIP), 0.1 μM each inner primer (F3 and B3), 0.2 μM each loop primer (FLoop and Bloop), 1 μl of template sample and 7.5 μl of Colorimetric LAMP Master Mix. The reaction was run in BioSan Dry block thermostat Bio TDB-100. The amplification products were detected by the reaction color shifted from red to yellow, which is based on the use of phenol red, a pH-sensitive indicator as instructed by the manufacturer. The products were also analyzed by electrophoresis on a 2% agarose gel when needed.

### Optimization of ASF colorimetric LAMP reactions

One ng of synthesized DNA template was used to perform the optimization experiment. Regarding identification of optimal incubation time, the LAMP reactions were incubated from 5 to 70 min at 60 °C. As for temperature optimization, the reaction mixtures were incubated at various temperatures from 50 °C to 70 °C for 30 min.

### LOD evaluation

The copy of synthetic DNA template was calculated using Endmemo program (http://endmemo.com/bio/dnacopynum.php; 1 ng is approximately equivalent to 3 × 10^19^ copies of the synthetic fragment of Topoisomerase II gene of ASFV). The synthesized DNA template was serially diluted to the indicated concentration and 1 μl of the diluted DNA was added to the LAMP reactions. Genomic DNA of ASFV was extracted from the known viral titer of 10^7^ HAD_50_/ml (50 percent haemadsorbing dose per ml) using the QIAamp DNA Mini Kit (Qiagen, USA). Extracted DNA was then serially 10-fold diluted and 1 μl of the diluted DNA was added to the LAMP reactions.

To identify the LOD value of LAMP assay when using crude serum sample, various amount of synthesized DNA template and extracted viral-genomic DNA were spiked into the serum collected from a healthy pig, producing the mimicked serum specimens containing the different concentrations of synthesized DNA template or extracted viral-genomic DNA. Next, the simulated serum samples was 10-fold diluted in the nuclease-free water and 1 μl of the diluted sample was added to the LAMP reactions.

### Specificity evaluation

Genomic RNAs of PRRSV (porcine reproductive and respiratory syndrome virus) and CSFV (classic swine fever virus) were extracted following the TRIzol method (TRIzol reagent, Invitrogen, California, USA). Extracted RNA of PRRSV was examined using VDx PRRSv qRT-PCR (NA/EU dual) Kit (Median Diagnostics Inc., Korea). Extracted RNA of CSFV was analyzed by agarose gel electrophoresis. The cDNAs of PRRSV and CSFV were then prepared using Maxime^™^ RT PreMix Kit (iNtRON Biotechnology Inc., Korea). The genomic DNA of ASFV (1 pg), genomic RNA (1 pg) and cDNA (1 pg) of PRRSV and CSFV were used to address the cross-reactivity of the LAMP kit.

To calculate the mismatch percentage of LAMP primers used among different ASFV strains, the complete genomes of various ASFV strains were downloaded from NCBI (https://www.ncbi.nlm.nih.gov/). The primers for LAMP were assembled to genomes of ASFV using Geneious Prime 2020.0.1 (https://www.geneious.com/) to calculate the number of different nucleotides. The mismatch percentage was defined by dividing the number of different nucleotides to total length of primers.

### Sensitivity evaluation

The sensitivity of ASF colorimetric LAMP assay was evaluated with extracted genomic-DNAs from 45 blood and serum samples of the pigs with clinical symptoms of ASF. Viral DNA was extracted by QIAamp DNA mini kit (Qiagen, USA) and 1 μl of extracted DNA was added to the LAMP reactions. The sensitivity of the LAMP kit was also determined with the direct use of 50 serum samples collected from the pigs in domestic farms. The serum samples were first 10-fold diluted in nuclease-free water and 1 μl of the diluted samples was applied directly to the LAMP reactions. For comparison, all 95 blood and serum samples were first applied to the QIAamp DNA mini kit (Qiagen, USA) for viral DNA extraction and then confirmed the ASFV presence by VDx^®^ASFV qPCR Kit (Median Diagnostics Inc., Korea) (Table S1).

## RESULT

### Optimization of ASF colorimetric LAMP reactions

The synthesized Topoisomerase II gene of ASFV was utilized as the template to carry out the LAMP reaction in the presence of the pH-sensitive indicator. The results indicated that the color change of the LAMP reactions corresponded to the amplified products generated (Fig. 1). Next, the optimal temperature and required time for the LAMP reaction for the detection of ASFV were defined as described in Materials and Methods section. The amplified product could be seen just after 10 minutes (min) of incubation by analyzing electrophoresis result. However, 15 min was the minimum incubation time to produce visualizable color change judged by eye (Fig. 2A). Besides, the amplicon was produced from 51 to 70 °C, however, the most distinctive color change was observed only from 55 °C (Fig. 2B). To ensure the outcome signal, the optimal reaction condition was set at 60 °C for 30 min. The LOD value, specificity and sensitivity of the developed colorimetric LAMP assay were thus identified at the optimized temperature and incubation time.

**Fig. 1.**
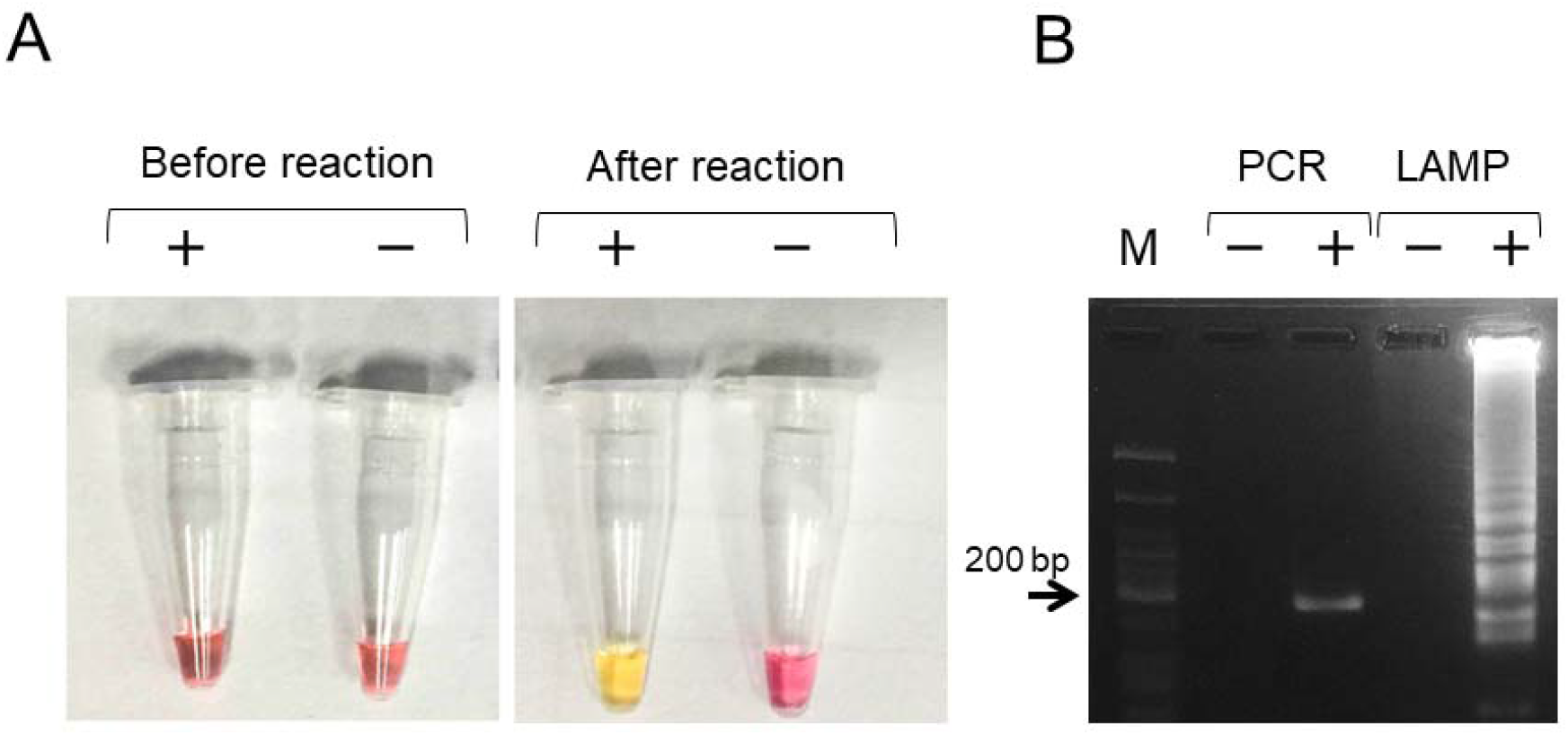
ASF colorimetric LAMP reaction using the synthesized DNA template. A) The amplified product was visualized by the color change from red to yellow (positive signal) and pink (negative signal). One ng of synthesized DNA template was used. The reactions were incubated at 60 °C for 30 min. B) LAMP product was analyzed by agarose gel electrophoresis and compared with the routine PCR result. Abbreviation, M: DNA ladder.

**Fig. 2.**
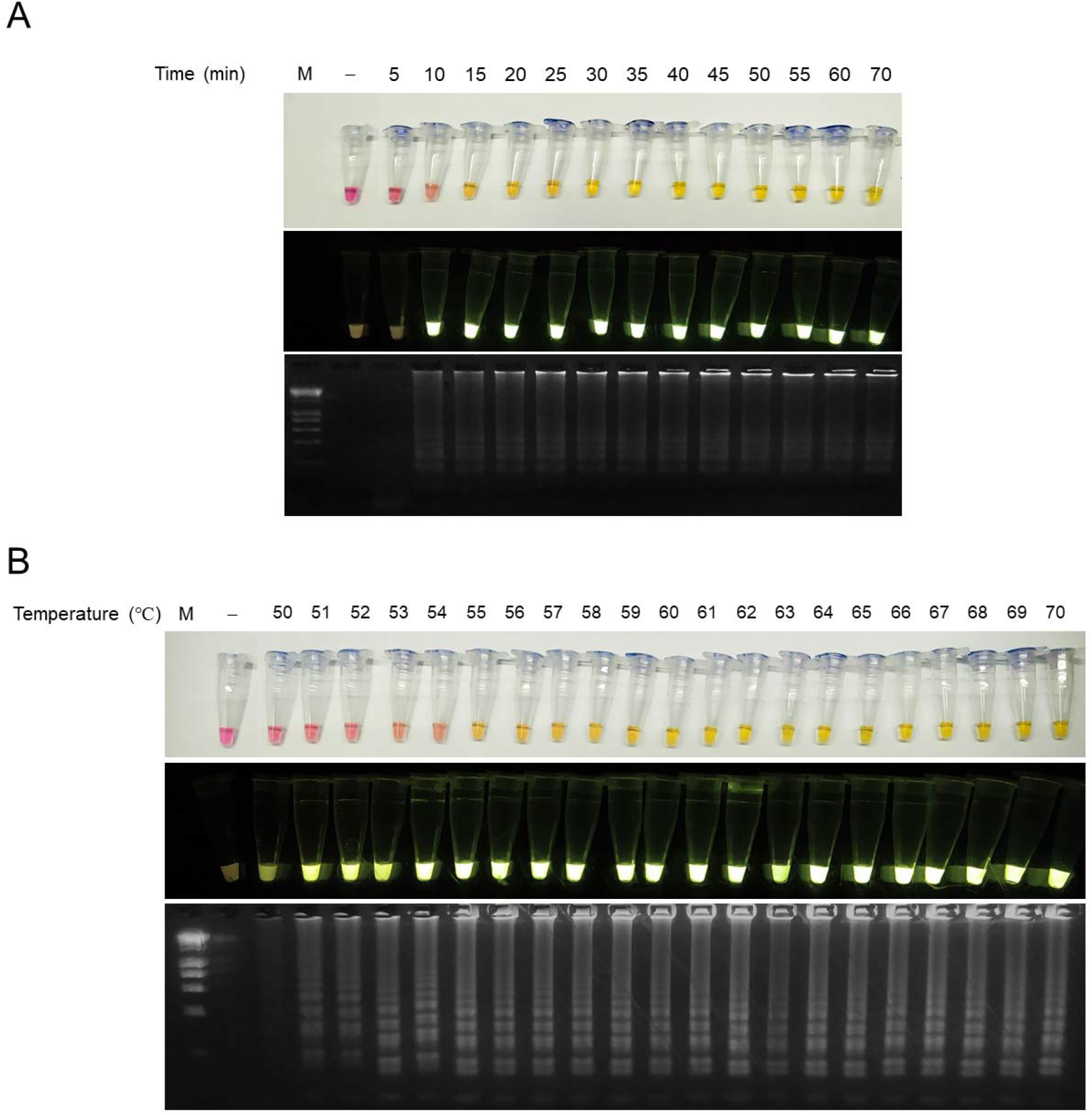
Optimization of the temperature and incubation time of the ASF colorimetric LAMP reaction. A) The reactions were incubated from 5 to 70 min at 60 °C. B) The reactions were incubated at different temperatures from 50 to 70 °C for 30 min. One ng of synthesized DNA template was used. Upper panel: LAMP results indicated by a pH-sensitive dye; middle panel: LAMP results indicated by adding SyBr Green to the reactions and visualized under UV light; lower panel: amplified products were analyzed by gel electrophoresis. Abbreviation, M: DNA ladder.

### LOD of ASF colorimetric LAMP assay

The LOD value of the LAMP reaction was evaluated using the synthesized DNA template and extracted viral-genomic DNA. As shown in Fig. 3A, roughly a single copy of the synthesized targeted gene per reaction was the lowest amount of synthesized DNA template that LAMP could detect. This LOD is identical to that of the commercial VDx^®^ ASFV qPCR Kit utilized in this study (Bokyu et al., 2019). Regarding viral genomic DNA detection, LAMP performance evaluated on ASFV’s DNA with known viral titer was estimated to be 1 HAD_50_/ml (Fig. 3B). The LOD of LAMP assay is thus approximately 14 times more sensitive than 10^1.16^ HAD_50_/ml of the commercial VDx^®^ ASFV qPCR Kit (Bokyu et al., 2019). Most importantly, using the simulated serum samples containing spiked DNA templates, LAMP reactions were successful at the lowest amount of 1 copy of the synthesized targeted gene and at the genomic DNA amount corresponding to 1 HAD_50_/ml of ASFV concentration. These values correspond to 10 copies of the targeted gene and 10 HAD_50_/ml of ASFV in serum samples (Fig. 3C and D). Note that from 10^5.3^ to 10^9.3^ HAD_50_/ml of ASFV could be detected in the blood of the infected pigs (McVicar, 1984). Hence, the LOD of ASF colorimetric LAMP assay is sufficient to identify ASFV in clinical samples.

**Fig. 3.**
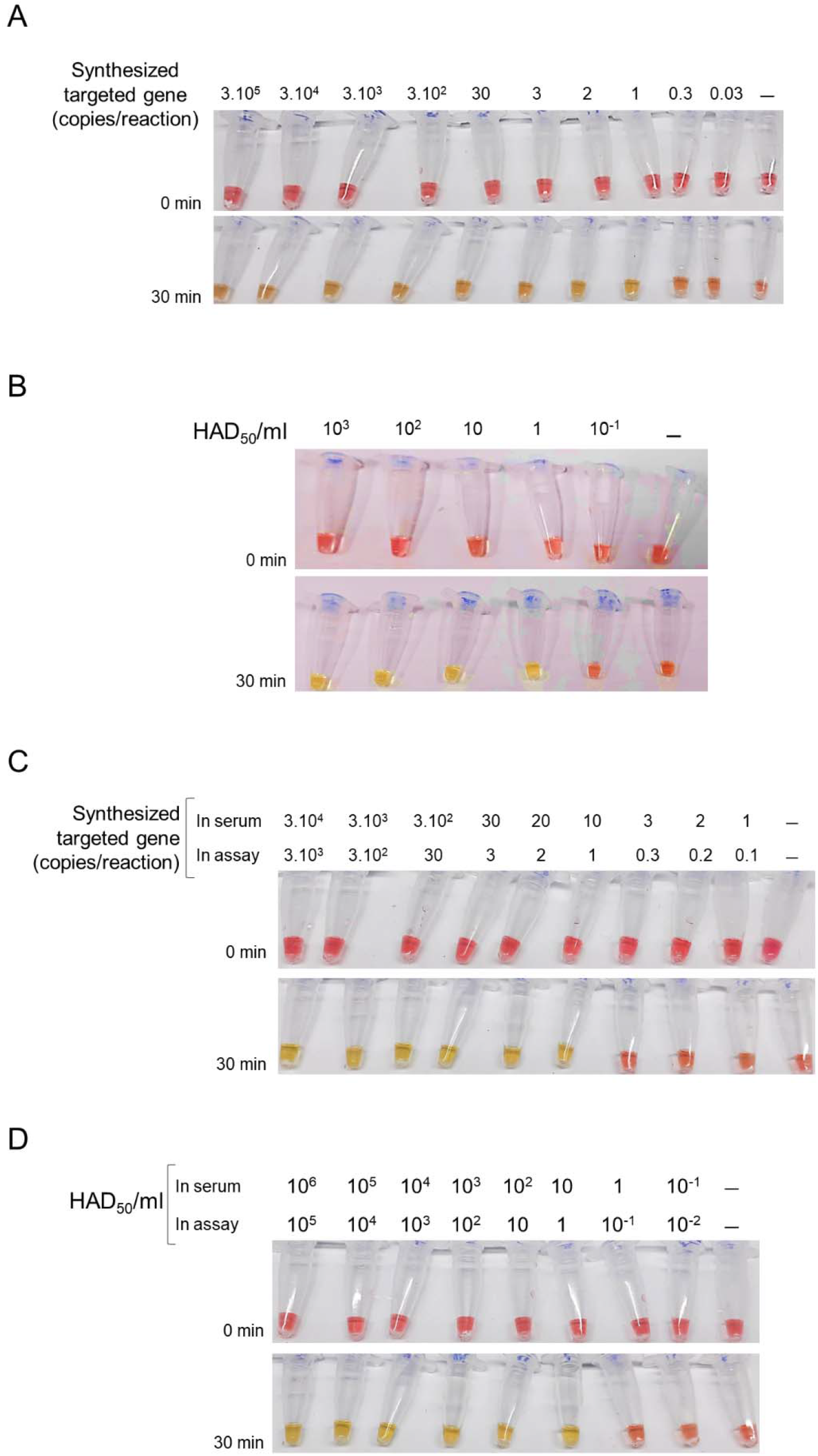
The LOD values of ASF colorimetric LAMP assay. LOD of LAMP assay was evaluated using (A) the synthesized DNA template and (B) the extracted viral genomic-DNA. LOD of LAMP assay was evaluated using the serum sample which was spiked with the defined concentrations of (C) the synthesized DNA template and (D) the extracted viral genomic-DNA. As for serum, the sample was 10-fold diluted in the nuclease-free water before added to the reaction. The amplified products were also confirmed by analyzed on gel electrophoresis.

### Specificity of ASF colorimetric LAMP assay

Previously, LAMP was demonstrated to have absolute specificity with no cross-reacting with CSFV was obtained (James et al., 2010). Consistently, the LAMP primer set used herein selectively detected the presence of ASFV DNA while no cross-reaction was observed with genomic RNAs of PRRSV or CSFV (Fig. 4A). Also, LAMP reactions produced negative signals for different bacteria DNAs tested including *Salmonella enterica, Staphylococcus aureus, Pseudomonas aeruginosa, Listeria monocytogenes* and *Vibrio parahaemolyticus* (data not shown). Thus, the primer set used is highly selective for ASFV.

**Fig. 4.**
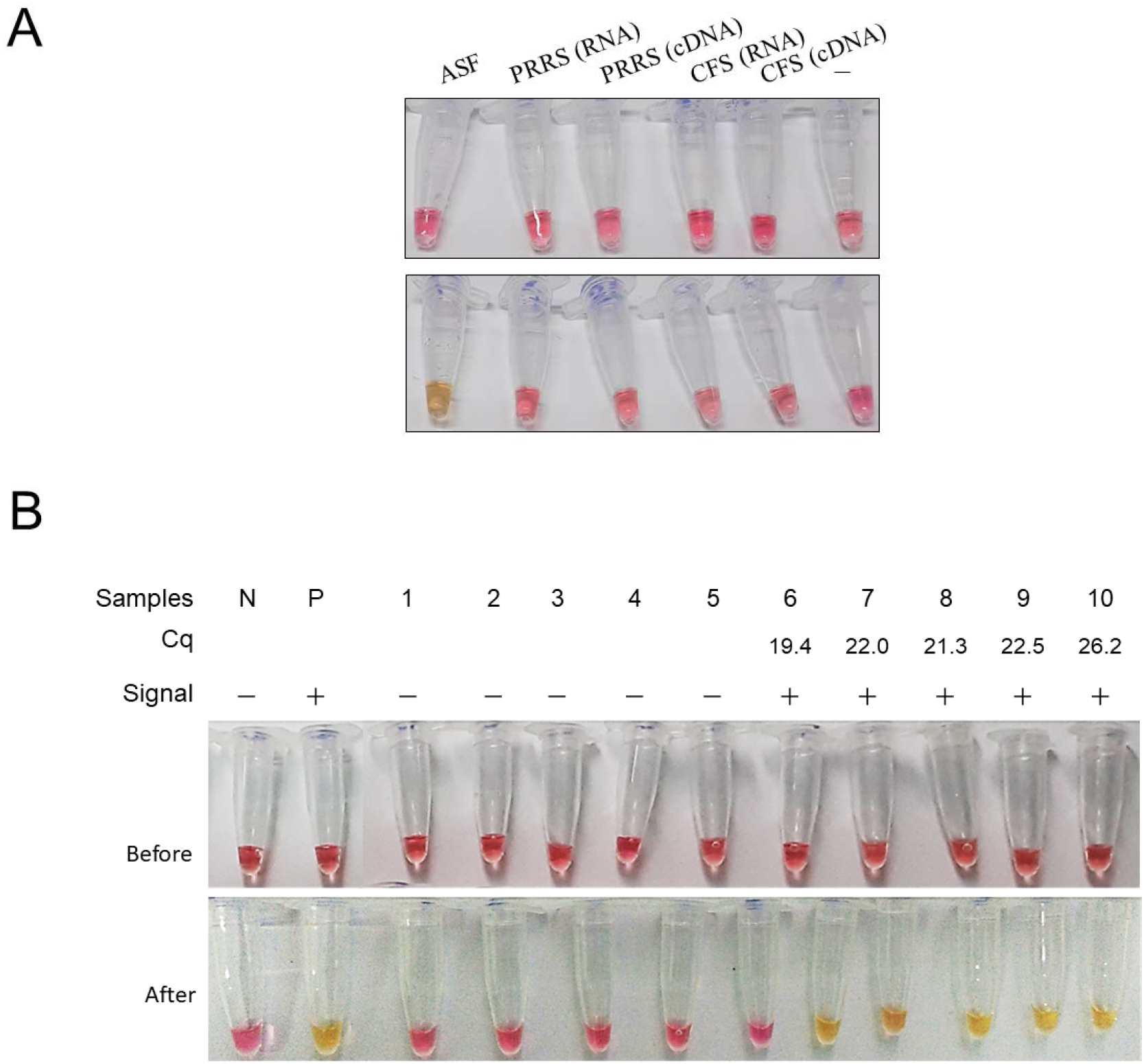
The specificity and sensitivity of ASF colorimetric LAMP assay. A) The specificity was evaluated with the extracted genomic RNAs and cDNAs of PRRSV and CFSV. (B) Representative results of LAMP assay using the extracted DNAs from 45 whole blood and serum samples. Abbreviation, N: negative control, P: positive control using the genomic DNA of VNUA/HY-ASF1 strain (Van Phan Le et al., 2019).

The ASFV strain found in Vietnam (VNUA/HY-ASF1) was shown to be 100% identical to China strain AnhuiXCGQ/China/2018 (GenBank accession no. MK128995) and some other genotype II strains of Europe, for example Georgia/2007/1 (FR682468.1), Estonia/2014 (LS478113), and Poland/2015 (MH681419) (Van Phan Le et al., 2019). Primer alignment shows that the mismatch rate between primer set used and DNA sequence of the ASFV strains mentioned above is only 2.55 percent while most of other ASFV strains examined exhibit the mismatch percentage of 2.55 % or lower (Table S2). Very few strains show the mismatch rate higher than 5%, and the highest percentage of 8.28% belongs to strain N10 (MH025919). The obtained data indicate that the primer set used is suitable for detection of ASFV strains in Vietnam, China, and most of countries in Europe and Africa.

### Sensitivity of ASF colorimetric LAMP assay

The sensitivity of LAMP assay was evaluated using extracted DNA of ASFV. Viral DNAs extracted from 45 whole blood and serum samples were examined both by real-time PCR and LAMP assay (Table S1, Fig. 4B). The results revealed that LAMP performance was identical to the commercial real-time PCR kit with 29 positive and 16 negative samples. The data indicate that the ASF colorimetric LAMP assay possesses the sensitivity and accuracy of 100% (Table 1).

**Table 1.** Sensitivity of ASF colorimetric LAMP assay evaluated using extracted genomic-DNA of ASFV

| VDx® ASFV qPCR Kit | LAMP |  | Total |
| --- | --- | --- | --- |
|  | Positive | Negative |  |
| Positive (n=29) | 29 | 0 | 29 |
| Negative (n=16) | 0 | 16 | 16 |
| Total (n=45) | 29 | 16 | 45 |
| Sensitivity: 100% |  | Specificity: 100% |  |

### Performance of ASF colorimetric LAMP assay with serum samples

ASF LAMP reactions were previously successful using crude samples, including whole blood and serum samples (Bredtmann, 2014). However, the use of whole blood and serum requires dilution, boiling step, and centrifugation for considerably better outcomes. Considering that whole blood contains a variety of components that could interfere with the amplification reaction while serum could be simply obtained by using a portable spin down machine, we utilized serum samples to perform LAMP assay. Unfortunately, direct use of 1 μl of serum sample in the reaction hindered the change of color although amplicon was still formed when analyzed by electrophoresis (data not shown). Hence, the serum sample was serially diluted with nuclease-free water and 1 μl of the diluted samples was then added to LAMP reaction mixture. Results show that the color change of reaction can be observed starting from 5-fold dilution (Fig. 5A).

**Fig. 5.**
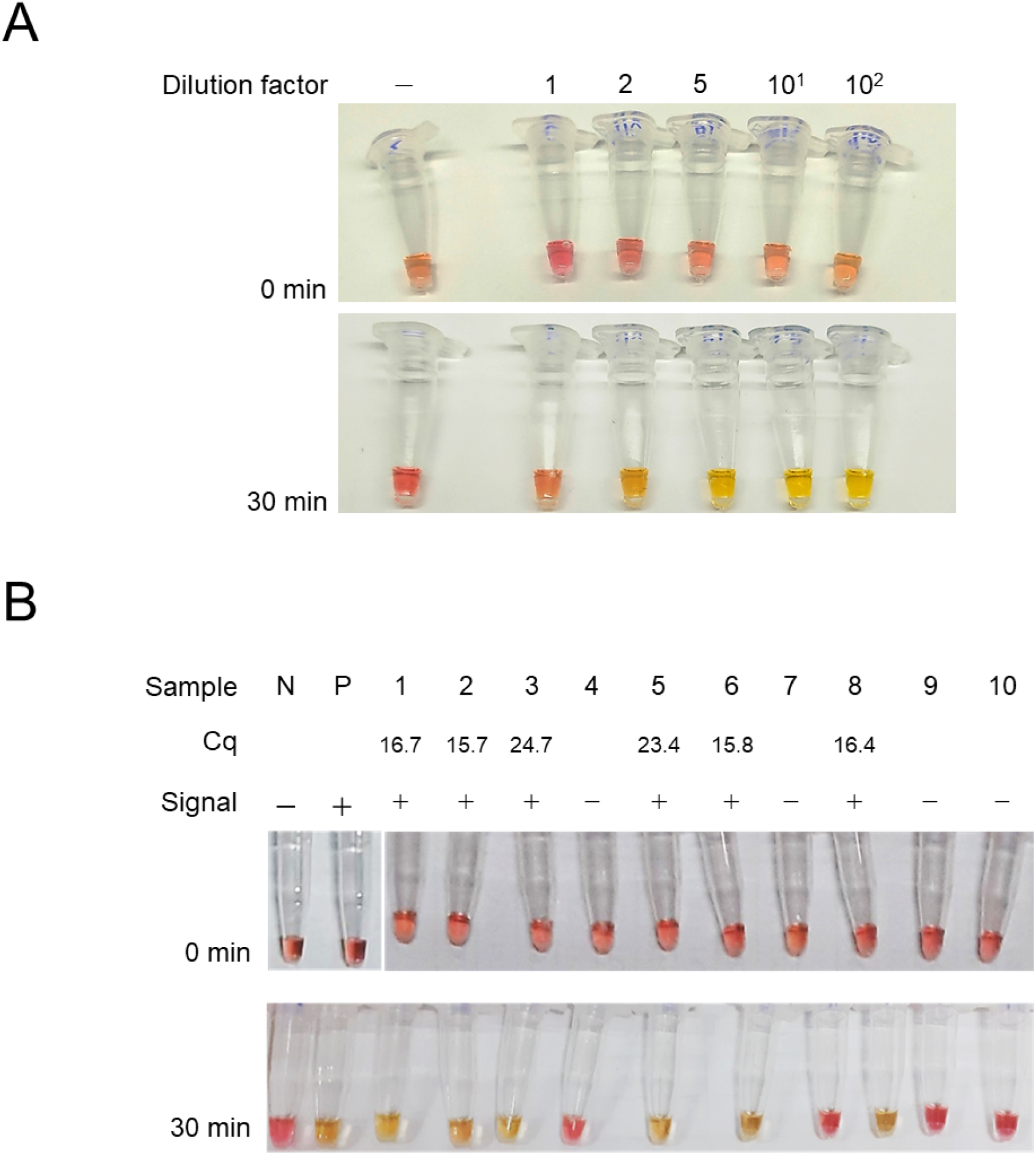
The sensitivity of ASF colorimetric LAMP assay using direct serum samples. (A) A serum sample containing ASFV was serially diluted in nuclease-free water to titrate the appropriate dilution rate for the LAMP reaction. (B) Representative results of LAMP assay using 50 serum samples directly. Crude serum samples were diluted 10-fold in nuclease-free water and 1 μl of the dilution was used for LAMP reactions. Abbreviation, N: negative control, P: positive control using the genomic DNA of VNUA/HY-ASF1 strain (Van Phan Le et al., 2019).

The performance of LAMP assay was subsequently evaluated with 10-fold dilution of 50 serum samples collected. Consistent with the results obtained with extracted genomic DNA, LAMP outcome was identical to the commercial real-time PCR kit with 35 positive and 15 negative samples, showing the sensitivity and accuracy of 100% (Table 2, Fig. 5B). Furthermore, the colorimetric LAMP assay performed well using the samples that were collected and stored for several months. It is worthy to mention that the direct use of serum samples successfully detected ASFV presenting at extremely low concentrations such as sample No 43 and 45 with Cq values of 33.66 and 33.0, respectively (Table S1). Note that a real-time PCR negative result is defined when Cq value is ≥35.

**Table 2.** Sensitivity of ASF colorimetric LAMP assay evaluated using crude serum samples

| VDx® ASFV qPCR Kit | LAMP |  | Total |
| --- | --- | --- | --- |
|  | Positive | Negative |  |
| Positive (n=35) | 35 | 0 | 35 |
| Negative (n=15) | 0 | 15 | 15 |
| Total (n=50) | 35 | 15 | 50 |
| Sensitivity: 100% |  | Specificity: 100% |  |

For application at the field, lyophilized reagents which are ready-to-use and does not require strict storage condition of low temperature would be considered. Thus, we attempted to investigate the LAMP performance with the lyophilized reagents. The reaction mixture containing the enzyme, primers, and dye was prepared and then freeze-dried. As expected, the LOD of freeze-dried LAMP reactions was maintained at 1 copy of the synthesized targeted gene per reaction and 1 HAD_50_/ml of ASFV (Fig. S1A and B). The specificity and sensitivity of detecting DNA were kept at 100% as well (data not shown). However, direct detection of serum samples was not successful. It could be explained by the more susceptibility of lyophilized reagents toward inhibitory components presenting in the crude serum, hindering the amplification process. Therefore, further investigation to select a factor(s) that could protect and stabilize the reaction components prepared for the freeze-drying process is strongly necessary. Also, the stability and performance of lyophilized assay under varied storage conditions is a subject that should be studied further.

## DISCUSSION

Nowadays, for better field-based screening and surveillance of infectious diseases, sensitive, accurate and low-cost diagnostic methods that do not require to be performed at centralized laboratories have been developed. Real-time PCR has been widely used for the accurate detection of animals carrying ASFV. However, this method always requires well-trained personnel and high-cost devices that is only feasible in the laboratory conditions and is not suitable for the screening and the fast decision-making process at the field. In contrast, the operational simplicity and isothermal setup make LAMP more feasible for the in-field diagnosis. Moreover, with the use of two or three specific primer sets that are complementary to different positions in the targeted region, the LAMP reaction could achieve extremely high performance in terms of sensitivity and specificity. Accordingly, the LAMP assay was established for the detection of ASFV previously. James et al succeeded to utilize LAMP to rapidly identify ASFV by targeting the Topoisomerase II gene and showed the LOD value of approximately 330 genome copies (James et al., 2010). However, the assay performance was just evaluated using extracted genomic DNA of the virus (James et al., 2010). Wu’s group used LAMP to detect ASFV successfully with the LOD of approximately 6 copies of the targeted gene per μl (Wu et al., 2016). Nevertheless, the reaction was carried out using only a synthesized template (Wu et al., 2016). Atuhaire and colleagues extensively evaluated the performance of LAMP and compared with the conventional OIE-recommended diagnostic PCR for detecting ASFV (Atuhaire et al., 2014). However, the study merely analyzed the assay performance using extracted viral genomic-DNA from blood and tissue samples (Atuhaire et al., 2014). Notably, Bredtmann *et al*. successfully analyzed LAMP performance to identify ASFV in pigs in Uganda using clinical crude samples, including whole blood, serum and tissue, without DNA extraction (Bredtmann, 2014). The results revealed that whole blood and serum could be applied directly to the LAMP assay. However, sample dilution was needed while boiling and centrifugation steps were remarkably beneficial for the reactions (Bredtmann, 2014). Also, even though this study developed LAMP to directly detect clinical specimens, the experimental setups with temperature changes and fluorescence-based detection method are not well suited for the in-field diagnosis.

Rather than conventional agarose electrophoresis, LAMP-amplified products could be visualized by the use of lateral flow strip (James et al., 2010) or SyBr Green (Atuhaire et al., 2014), a dye only fluoresces upon binding to double-stranded DNA molecules when exposed to a UV light source. Also, the reaction reading can be performed similarly to real-time PCR with Melting Curve Analysis or the inverse protocol called Re-association Curve Analysis (Bredtmann, 2014). Regarding feasible application at the field, using the suitable dye to instantly read out the reaction results is more appropriate. However, the use of SyBr Green was reported to be very sensitive to contamination in the reaction mixture and could produce false-positive results (Atuhaire et al., 2014). Alternatively, recently, advantaging on the pH reduction of LAMP reaction based on the production of protons that occurs from the extensive DNA polymerase activity, a pH-sensitive dye which does not interfere with the reaction was introduced to indicate the outcome signal (Tanner, Zhang, & Evans, 2015). The color change due to pH decrease could be simply read by naked eyes (Tanner et al., 2015). Thus, colorimetric detection allows for the amplified product to be detected in a timely manner, without the demand for extended work and specialized apparatus. Moreover, a pH-sensitive dye is considerably cheaper than SyBr Green, which is more advantageous to be developed as an in-field diagnostic test kit. Accordingly, very recently, Wang et al introduced a real-time LAMP and visual LAMP that utilized the pH-sensitive dye for early diagnosis of ASF with a detection limit of 30 copies per μl of pUC57 containing p10 gene sequence (Wang et al., 2020). The visual LAMP assay was comparable to the well-established real-time PCR (Wang et al., 2020). However, the assay was evaluated with extracted DNA that limited it application at the field.

Due to the lack of evaluation on crude samples or simple visualization of reaction outcome, earlier developed ASF LAMP assays were still less feasible for field applications with restricted resources. The purpose of this study was to apply and evaluate a current developed colorimetric LAMP assay to rapidly and accurately detect ASFV in pigs grown in Vietnam as an alternative method to real-time PCR for diagnostic purposes at the field. The optimal conditions for ASFV detection by colorimetric LAMP reaction were identified as 60 °C for 30 min, even that the amplicon could be observed as early as after 15 min of incubation. Because the testing time is critically important for the in-field diagnosis of infection, the fact that the results are clearly visible to naked eyes after 15-30 min makes this colorimetric LAMP assay an ideal option to detect ASFV at the field. Moreover, the ASF colorimetric LAMP assay was extremely sensitive for the detection of ASFV, comparable to the commercial real-time PCR kit. Also, no cross-reactivity was detected with the closely related viruses and several bacterial genomic-DNAs, indicating that the developed LAMP assay is highly specific, in agreement with the previous study (James et al., 2010). Importantly, the sensitivity and LOD of colorimetric LAMP assay were not affected by the use of crude clinical samples that were not go through the DNA extraction process, significantly reducing the time and cost of the assay, making LAMP among the most effective candidates for the in-field detection of ASFV.

## CONCLUSION

This is the first study applying and evaluating the colorimetric LAMP assay using field samples in the detection of ASFV from infected domestic pigs in Vietnam. The colorimetric LAMP assay was shown to be as sensitive as the recommended real-time PCR assay in detecting ASFV. Moreover, unlike real-time PCR, the best advantage of LAMP is the possibility of working directly on crude clinical samples, without the need of a prior DNA extraction step. Hence, the developed colorimetric LAMP assay is a highly specific, sensitive, rapid and simple alternative approach to real-time PCR for the precise and quick detection of ASFV at the field with limited testing facilities. Also, to commercialize the ASF colorimetric LAMP kit, further evaluations of the performance of LAMP assay in long-term and storage conditions should be addressed.

## ETHICS STATEMENT

The authors confirm that the ethical policies of the journal, as noted on the journal’s author guidelines page, have been adhered to and the appropriate ethical review committee approval has been received. The Vietnam Veterinary Association’s guidelines for the Care and Use of Laboratory Animals were followed.

## ACKNOWLEDGEMENT

This work was supported by the Vietnam National Project under the Project Code No: DTDL.CN-53/19 and by NTTU Foundation for Science and Technology Development under grant number 2020.01.020.

## DATA AVAILABILITY STATEMENT

The data that support the findings of this study are available from the corresponding author upon reasonable request.

