## Supplementary material for "Direct colorimetric LAMP assay for in-field detection of African swine fever virus: a validation study during an outbreak in Vietnam": Figure S1

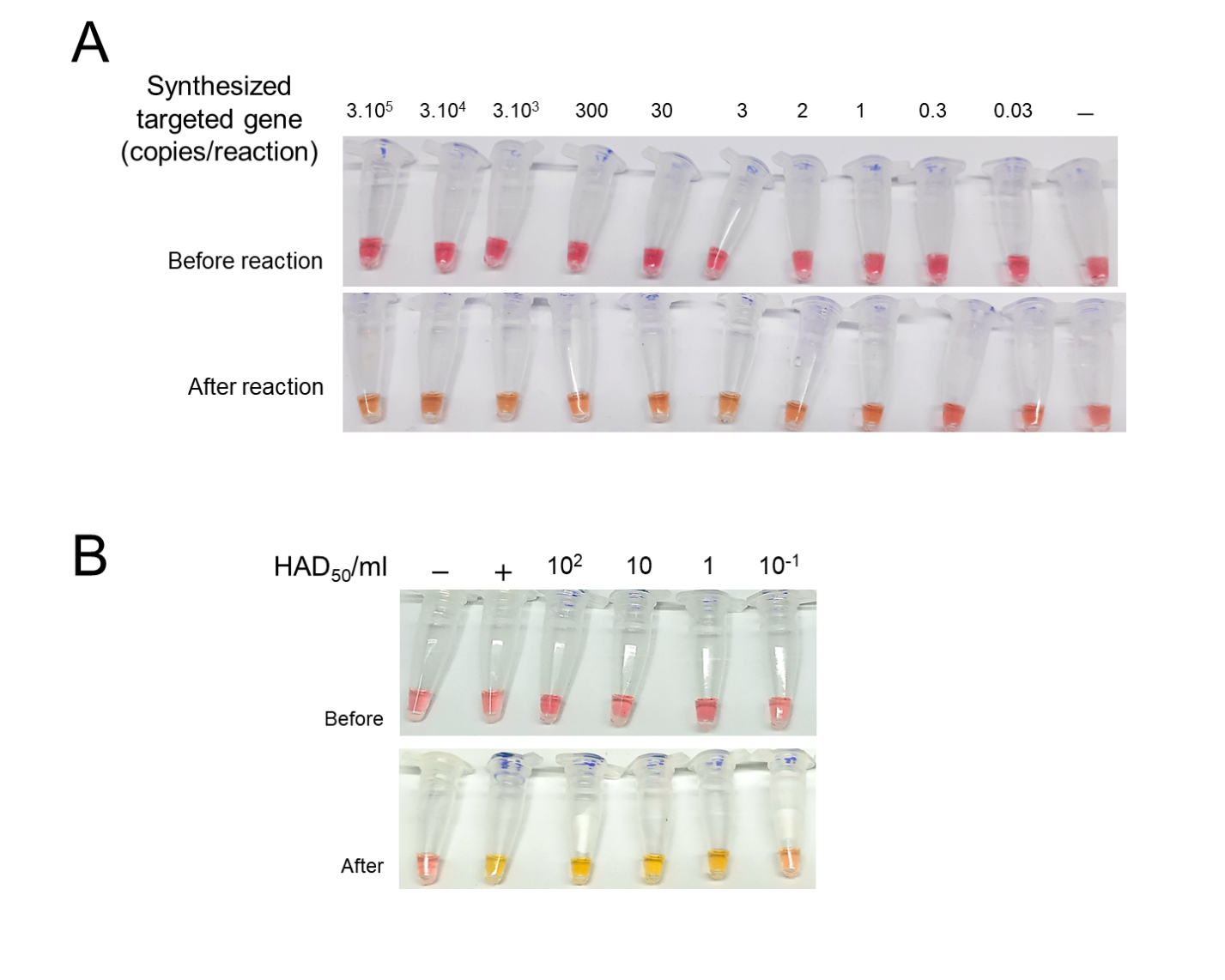

**Figure S1. The performance of colorimetric LAMP dried reagents.** A) The LOD of the assay was evaluated using synthesized DNA template. B) The LOD of the assay was evaluated using the extracted genomic DNA.
